# HistMapR: Rapid digitization of historical land-use maps in R

**DOI:** 10.1101/109504

**Authors:** Alistair G. Auffret, Adam Kimberley, Jan Plue, Helle Skånes, Simon Jakobsson, Emelie Waldén, Marika Wennbom, Heather Wood, James M. Bullock, Sara A. O. Cousins, Mira Gartz, Danny A. P. Hooftman, Louise Tränk

## Abstract

**1.** Habitat destruction and degradation represent serious threats to biodiversity, and quantification of land-use change over time is important for understanding the consequences of these changes to organisms and ecosystem service provision.

**2.** Comparing land use between maps from different time periods allows estimation of the magnitude of habitat change in an area. However, digitizing historical maps manually is time-consuming and analyses of change are usually carried out at small spatial extents or at low resolutions.

**3.** We developed a method to semi-automatically digitize historical land-use maps using the R environment. We created a number of functions that use the existing *raster* package to classify land use according to a map’s colours, as defined by the RGB channels of the raster image. The method was tested on three different types of historical land-use map and results were compared to manual digitisations.

**4.** Our method is fast, and agreement with manually-digitised maps of around 80-92% meets common targets for image classification. We hope that the ability to quickly classify large areas of historical land-use will promote the inclusion of land-use change into analyses of biodiversity, species distributions and ecosystem services.

## Introduction

Historical land-use maps represent valuable sources of information in ecology. In addition to the estimation of land-use change over time (Skånes & Bunce 1997; Swetnam 2007), historical map data are commonly coupled with species observations to relate land-use change to changes in biodiversity (Saar *et al.* 2012; Cousins *et al.* 2015; Hooftman, Edwards & Bullock 2016) and ecosystem services over time (Jiang, Bullock & Hooftman 2013; Willcock *et al.* 2016).

At present, most studies involving the analysis of historical land-use are carried out at landscape scales (Swetnam 2007; Cousins 2009; Saar *et al.* 2012), while analyses at larger spatial scales are uncommon (Hooftman & Bullock 2012; Cousins *et al.* 2015; Willcock *et al.* 2016). This is because digitization of historical land-use maps most commonly involves the time-consuming manual delineation of different land-cover types on scanned, georeferenced historical maps using a desktop GIS or illustration program. As a result, historical land-use (change) rarely features in analyses of biodiversity and species distributions following environmental change at large spatial scales (Hill *et al.* 2002; Powney *et al.* 2014), despite the acknowledgment of land-use change as the principal determinant of biodiversity loss worldwide (Newbold *et al.* 2016)

The *HistMapR* package contains a set of functions that allow a fast and accurate digitization of historical land-use maps in R (R Development Core Team 2015). Map colours are defined by the combination of values (0-255) of the RGB (Red, Green and Blue) channels of a raster image. Calling functions from the *raster* package (Hijmans 2016), our method uses RGB values to classify land use according to user-defined colours. We describe the method, before demonstrating it using three historical map series and comparing outputs to manual digitizations.

## Materials and methods

### Classification method

#### Step 1. Image smoothing (function: *smooth_map*)

Scanned historical paper maps contain inconsistencies in colour due to variations in map production, age and the quality of scanning. The first function applies a Gaussian smoothing to the input raster, calling the *focal* function from the *raster* package. Each pixel in each RGB channel is assigned the mean value from a user-defined window of *n* pixels surrounding the target pixel. In addition, RGB values below a user-defined threshold can be removed. This allows small patches of dark colour, for example denoting place names and property boundaries to be ‘smoothed over’ so that they do not interfere with the land-use classification. Finally, the smoothed raster is cut to the dimensions of the input raster to remove the halo effect, which occurs where the smoothing process spreads pixel values into any non-image areas of the raster.

#### Step 2. Assign user-defined colours (function: *click_sample*)

This function requires the user to define the colours for each land-use category from the smoothed map, calling the *raster* package's *click* function. Clicking a number of times within each category from across the image ensures that the full range of colour tones is sampled. A colour table containing maximum, median and minimum RGB values for each category is produced, and the associated colours can be plotted for inspection with the extra function *plot_colour_table*, calling functions from the packages *gridExtra* (Auguie 2016) and *ggplot2* (Wickham 2009).

#### Step 3. Test classification and write to raster file (function: *class_map*)

Each pixel in the smoothed raster is then assigned to a land-use category according to the colour table produced in the previous step (so-called parallelpiped classification). Categories are assigned from the first row of the colour table and down, meaning that if a pixel contains RGB values falling within the range of several categories, it is to the nethermost category in the table that the pixel is assigned in the final classification. Rearranging the colour table allows the user to choose which categories should take precedence over others in the case of overlap. Additionally, the range of RGB values in each category can be expanded by a chosen number of standard errors to account for the likelihood that the most extreme RGB values for each category were not clicked in the previous step. The effects of various standard errors can be visually inspected using *plot_colour_table.* Finally, the RGB values of some pixels are likely to fall outside all categories in the colour table. These exceptions can be assigned to an existing category or left unclassified. The effects of rearranging the colour table, assigning error values and exception categories can be assessed by plotting within the R environment and by writing to a raster file for examination in a GIS program.

#### Additional steps

Different maps from the same series may require different colour table arrangements to achieve optimal results. In such cases maps must be reclassified so that raster categories match among maps prior to analysis and joining maps together to cover larger areas. Additionally, in two of the three historical map series below, surface water was not denoted in a way which meant that they could be adequately classified as a separate land-use category using their RGB values. In these cases we used the function *gdal_rasterize* in the package *gdalUtils* (Greenberg & Mattiuzzi 2015) to burn a modern water vector layer onto the digitized raster. The *HistMapR* package and documentation are hosted at <u>https://github.com/AGAuffret/HistMapR/</u>. Detailed example scripts and input maps are available on Figshare (Auffret *et al.* 2017).

### Case study examples

#### Dorset, UK - The Land Utilisation Survey of Great Britain (1930s)

The UK Land Utilisation Survey was led by Stamp (1931). Sheet 140 over Weymouth and Dorchester covers was 800 km^2^ and was mapped at the 1:63 360 scale, depicting the following land-use types: [1] forest and woodland, [2] arable land, [3] meadow and permanent grass, [4] heath and moorland, [5] gardens, orchards and allotments, [6] urban and industrial areas, [7] inland water (**Figure 1a**). We used *HistMapR* to classify land use on this map into a raster file containing these seven categories at a resolution of 8 m^2^. Total computing time was around 30 minutes on a standard computer, excluding time spent assigning colours with *click_sample* and testing. We then compared our digitized map to Hooftman & Bullock (2012), who manually-digitized maps over the county of Dorset. To be able to compare outputs we first reclassified land-use categories, merging [5] & [6] to match the manual digitization. The manual digitization was rasterized using *gdal_rasterize*, and both digitizations were aggregated by a factor of five to try to reduce the effect of differences in georeferencing, before being masked by each other using *raster*'s *mask* function to ensure that they had the same extent. Total agreement between the two digitizations was calculated by identifying the fraction of corresponding pixels that were classified into the same category. We also calculated the fraction of pixels assigned to each map category in the manually-digitized map the fraction of pixels that were categorized as each category in the HistMapRdigitization. Finally, the total fraction assigned to each category in each digitization, and the root-mean-square deviation (RMSD) of cover between digitizations was calculated.

#### Södermanland, Sweden - District Economic map of Sweden (1859-1934)

This map series (AKA The Hundred map; Swedish: *Häradsekonomiska kartan*) describes major land use, settlements and infrastructure (**Figure 1b**). We digitized 11 maps in the county of Södermanland (scale 1:20 000) that were manually digitized by Cousins *et al.* (2015). Each map covers approximately 105 km^2^, and the manual digitization classified land use into nine categories. We classified land use into the general categories of forest, arable land, meadow/dwelling and water (using a modern layer, see above) at a 4 m^2^ resolution. Computing time was approximately 15 minutes per map. Comparison of the *HistMapR* and the manual digitization was performed as above for each map sheet, with RMSD also calculated for each category individually.

**Figure 1.**
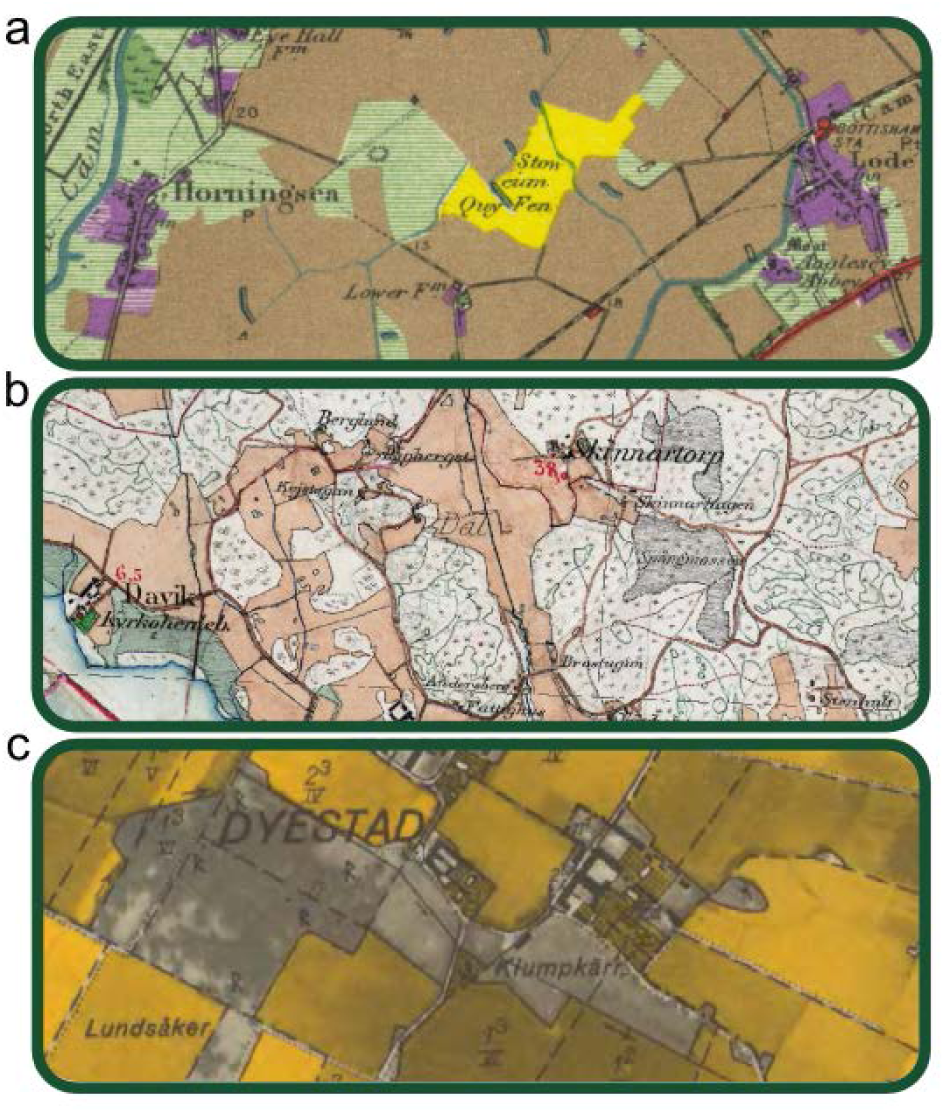
*Illustrative examples from (a) the Land Utilisation Survey of Great Britain, (b) the Swedish District Economic map and (c) the Swedish Economic map. Map images copyright (a) Audrey N. Clark, (bc) the Swedish Agency Lantmäteriet.*

#### Southern Sweden - The Economic map of Sweden (1935-1978)

The Economic map series (*Ekonomiska kartan*) was successor to the District Economic Map, published 1935-1978 and covering the whole of Sweden. In southern Sweden, each sheet covers 25 km^2^ at the 1:10 000 scale. The maps consist of a monochrome aerial orthomosaic, with arable land, gardens and pasture on former arable fields coloured yellow, and additional information such as roads, larger buildings and boundaries in black (**Figure 1c**). We classified 7069 maps from the 15 southernmost counties in Sweden, corresponding to an area of 176 725 km^2^, at a 1 m^2^ resolution.

Maps were split according to county, and then visually inspected and split into a number of groups using a file manager or GIS program according to the relative colour tones present in the map. For most counties, this resulted in 5-20 groups containing anything from a few up to 329 maps. Within each group, a representative map was digitized using *HistMapR* into arable land etc. (yellow), forest (darker shades - trees present in the map image) and other open land (lighter shades – no trees). Classification settings of the selected map were tested on another map within the same group before running the method in a for-loop or computer cluster to digitize all maps in the group unsupervised. Computation time was 5-10 minutes per sheet. These batched classifications were inspected in a GIS program and groups or individual maps re-run with different settings as needed. Water was added using a modern vector layer as described above. For verification, we took 34 manually-digitized maps from across the study region, 0.79-139 km^2^ in area. These were either digitizations of the Economic maps themselves (Gartz 2015; J. Plue unpublished data), or stereographic interpretations of contemporary aerial photographs (Skånes & Bunce 1997; Cousins & Eriksson 2008; Cousins 2009). Land-use categories were changed to match our classification and comparisons were carried out as described above.

## Results

We found *HistMapR* to be both fast and straightforward. Comparing outputs with manual digitizations showed good agreement for all three map series. The map over Dorset showed a 92% overall agreement at the pixel level, with the majority of pixels in each land-use category being classified to the same category across digitizations (**Figure 2a**). Agreement in the Swedish map series was generally around 80-90% **(Fig 2b-c)**, with pixels again mostly classified into the same categories using both methods. In the Economic map, there was less agreement for open areas compared to arable and forested land.

**Figure 2.**
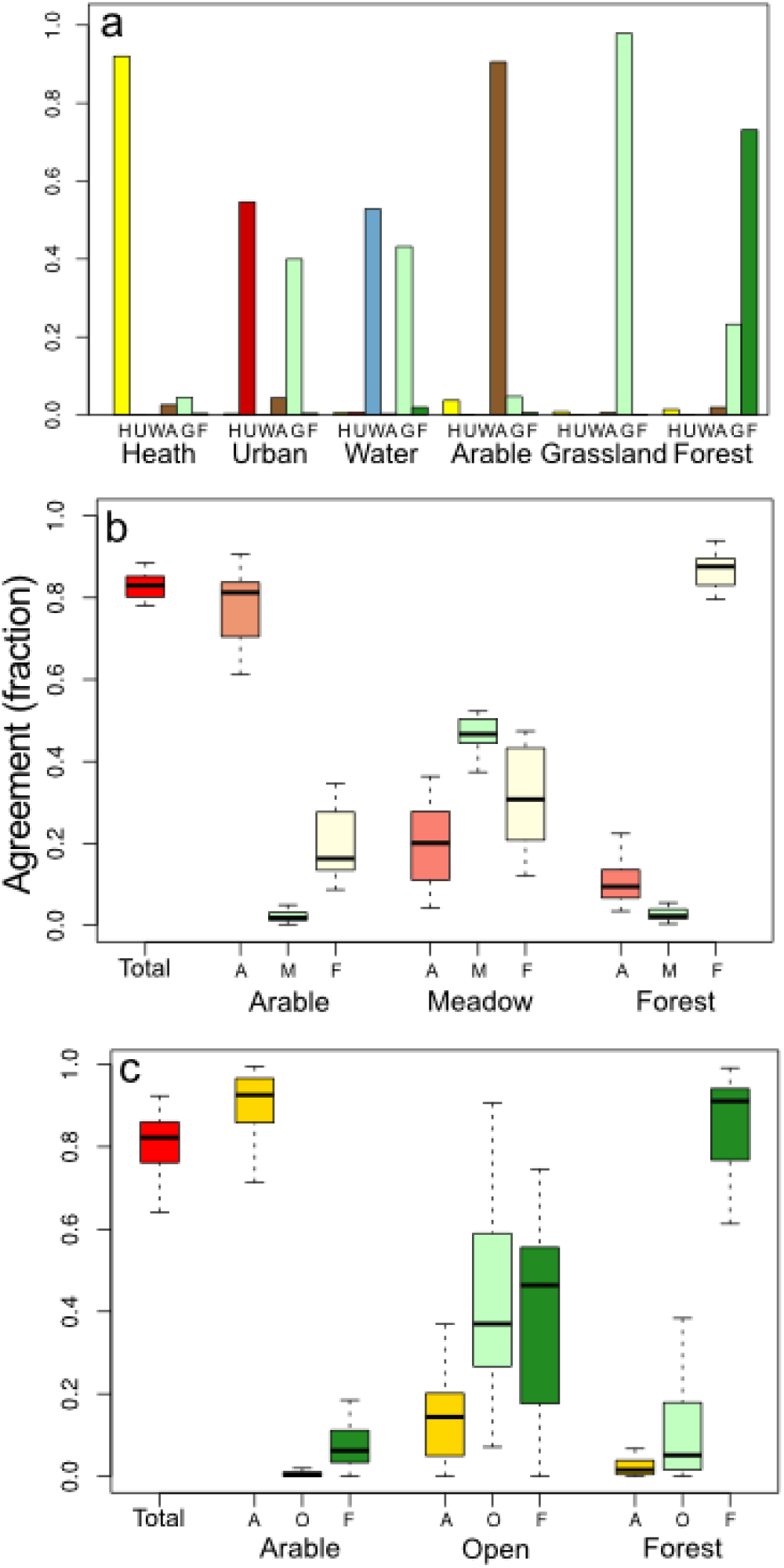
*Fraction of pixels assigned to the same land-use category from manual and HistMapR digitizations, and the fraction of pixels in each manual digitisation that are assigned to each map category in the HistMapR digitisation of (a) the Land Utilisation Survey of Great Britain (1 map), (b) the Swedish District Economic map series (11 maps) and (c) the Swedish Economic map series (34 manual digitizations). Boxes represent upper and lower quartiles, thick lines show the median, and whiskers the dataset range without outliers (observations falling outside the quartiles +/- 1.5 **×** the interquartile range). Colours match original map shadings.*

**Figure 3.**
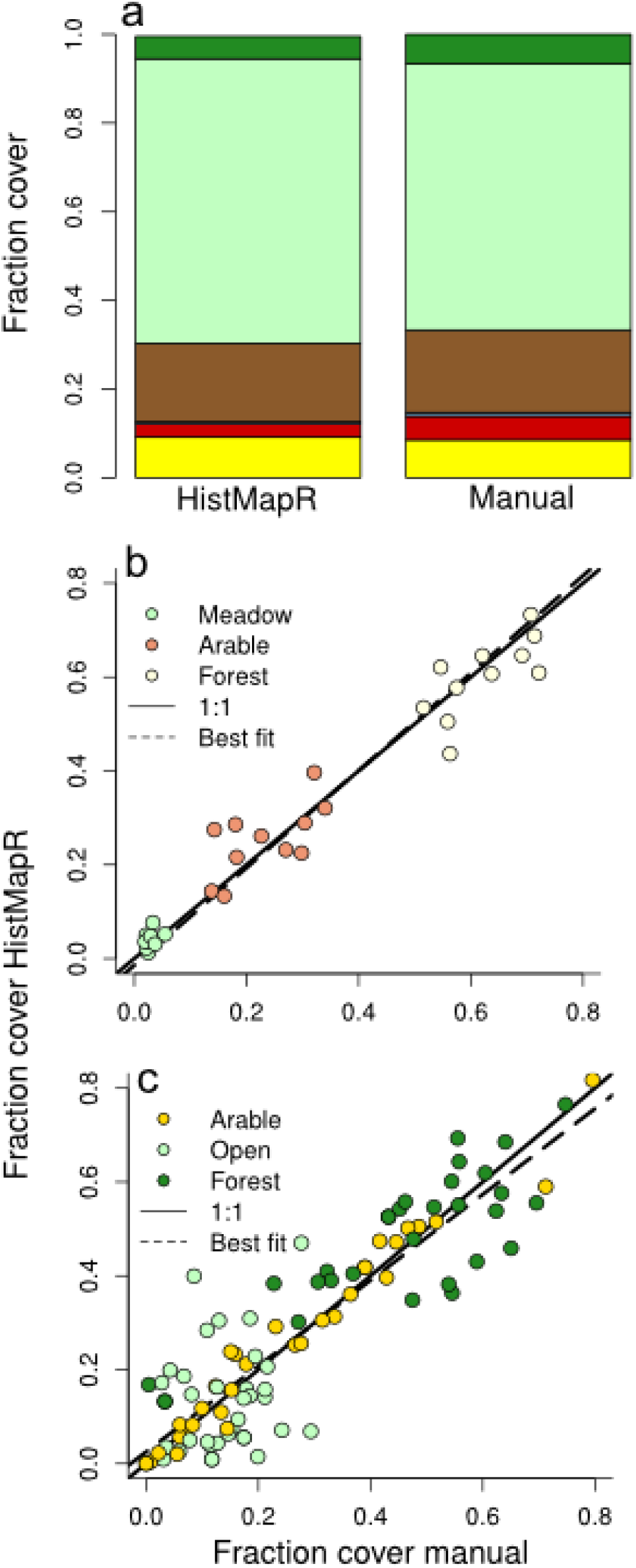
*Fraction of land cover assigned to each land-use category in (a) the Land Utilisation Survey of Great Britain (1 map), (b) the Swedish District Economic map series (11 maps) and (c) the Swedish Economic map series (34 manual digitizations). Colours match original map shadings.*

Overall share of land-use categories was very similar between the *HistMapR* and manual digitizations. In Dorset, deviation (RMSD) across categories for the whole map was 2%, with grassland over-represented at the expense of small proportions of most other categories **(Figure 3a)**. For the District Economic Map, deviation was 4.6% for all categories combined, with values of 2%, 6%, and 6% for arable, meadow and forest categories respectively (**Figure 3b**). Deviation in the Economic map was 9% for all categories, 4% for arable, 12% for open land and 12% for forest (**Figure 3c**).

## Discussion

We have developed a method for a rapid and accurate semi-automated classification of historical land-use using open-source software. Pixel-level agreement between *HistMapR* and manual digitizations was high for all map series **(Fig 2a-c)**, meeting commonly-set targets for land-cover classification accuracy (Foody 2002). Deviation of fractional cover of land-use categories between digitizations was usually within a few percent, both at the overall and category level (**Fig 3a-c**), while time savings were significant. We estimate that the manual digitisation of our classified area of the Dorset map (Hooftman & Bullock 2012) took approximately 3-4 weeks to complete, including familiarization with the area and partial method development, compared to the 30-minute *HistMapR* digitization. The almost 1700 km^2^ study area of the District Economic Map took around two months to manually digitize for Cousins *et al.* (2015) compared to 1-2 days’ work using our method.

Despite good results, there were sources of error, which differed between map series. On the smaller-scale UK map, land-use categories that largely consisted of small and linear elements, such as urban areas (including roads) and surface water were more affected by the smoothing function. This meant that the boundary areas of these land-use types differed in colour from core areas and therefore were often classed according to the exceptions argument, in this case grassland (**Figure 2a**). In the District Economic map series, disagreement at the pixel level arose due to map age and poor scan quality, resulting in variation within and between land-use categories in each map sheet (**Figure 2b**). The disagreement relating to forest and open land across digitization for the Economic maps (**Figure 2c**, **3c**) was largely due to the character of the map series. Only arable land, gardens and pasture on former arable fields were formally mapped, with other land-use types only visible as part of the underlying image. This means that all manual digitizations involved users to actively determine the level of tree-cover needed to discriminate parcels of wooded from open land. On the other hand, discrimination between relatively-darker and lighter colours (forest and open land, respectively) could only take place at the whole map level when using the *HistMapR* method, and pixels were then classified as such regardless of patch size. Furthermore, over one-third of 34 the manual digitizations used for comparison were based on corresponding aerial photographs rather than the Economic maps themselves, meaning that in several cases arable fields in the Economic maps were classified as open grassland in the manual digitization and vice versa, thus introducing an additional source of disagreement.

Our results show that despite some pixel-level disagreement, the resulting effect on relative cover of land-use types is generally low **(Figure 3)**. It is also important to point out that disagreement between *HistMapR* and manual digitizations does not equate to our maps being incorrect. The issue of delineating land-use categories is a problem for any land-cover classification. With *HistMapR*, users can tailor classification to suit their specific research questions and minimise other potential sources of error according to the historical map in question. Moreover, imperfect georeferencing between digitizations meaning that layers do not always perfectly overlap leads us to believe that actual agreement may even be higher. Our digitizations also represent raw outputs from the R environment, and the potential for improving small-scale accuracy with other GIS tools remains, while retaining significant time savings compared to manual digitization.

Although manual land-use classification results in a more accurate and detailed digital representation of historical maps, our method is highly useful for a range of applications in ecology. To efficiently classify broad land-use categories over large areas is extremely valuable for quantifying the magnitude of habitat loss over time. This could lead to a greater understanding of the anthropogenic drivers of changes in species diversity and distributions, enabling better predictions of future responses to change at multiple spatial scales.

## Acknowledgements

We are grateful to R. Hijmans for creating the *raster* package upon which our method heavily relies. For the Dorset map, scanned images were provided through <u>http://www.VisionofBritain.org.uk</u>, showing material from The Land Utilisation Survey of Great Britain, 1933-49, ©Audrey N. Clark. The manually-digitised map was taken from georeferenced scans of Dorset created by JMB and DAPH under DEFRA licence 10001880. Swedish maps ©Lantmäteriet made available to Stockholm University on licence I2014/00691. Many thanks go to the Swedish OpenStreetMap community for georeferencing the Economic Map. A. Smith and P. Platts gave useful help and advice. This work is funded by the Swedish research council Formas (2015-1065).

## Author contributions

AGA conceived the project. AGA and AK developed the method and created the functions. AGA, AK, SJ, JP, HS, EW, MW, HW tested the method and digitised maps. JMB, SAOC, DAPH, MG, JP, HS, LT manually digitised maps used for verification. AGA analysed the data and led the writing in close consultation with AK. All co-authors assisted with edits and approve publication.

## Data accessibility

### Code and example scripts

The *HistMapR* package and documentation are hosted at <u>https://github.com/AGAuffret/HistMapR/</u>. Detailed example scripts and input maps are available from Figshare <u>http://dx.doi.org/10.17045/sthlmuni.4649854</u> (Auffret *et al.* 2017).

### Maps

All Swedish District Economic and Economic maps that we digitized using our method are also available from Figshare for download and use, along with the manually-digitized maps used for verification (Auffret *et al.* 2017). The Dorset maps are under 3^rd^ party copyright.

Scanned Swedish historical maps can be found at <u>http://historiskakartor.lantmateriet.se/en</u> (Accessed: 2 February 2017). We used Lantmäteriet’s open-access terrain map for contemporary water layers, available from <u>https://www.lantmateriet.se/sv/Kartor-och-geografisk-information/Kartor/oppna-data/hamta-oppna-geodata/</u> (In Swedish; Accessed: 2 February 2017).

## References

Auffret, A.G., Kimberley, A., Plue, J., Skånes, H., Jakobsson, S., Waldén, E., Wennbom, M., Wood, H., Bullock, J.M., Cousins, S.A.O., Hooftman, D.A.P., Gartz, M. & Tränk, L. (2017) Data from: HistMapR: Rapid digitization of historical land-use maps in R. figshare data repository doi: 10.17045/sthlmuni.4649854.

Auguie, B. (2016) gridExtra: Miscellaneous Functions for “Grid” Graphics. R package version 2.2.1, url: http://CRAN.R-project.org/package=gridExtra.

Cousins, S.A.O. (2009) Landscape history and soil properties affect grassland decline and plant species richness in rural landscapes. Biological Conservation, 142, 2752–2758.

Cousins, S.A.O., Auffret, A.G., Lindgren, J. & Tränk, L. (2015) Regional-scale land-cover change during the 20th century and its consequences for biodiversity. AMBIO, 44, 17–27.

Cousins, S.A.O. & Eriksson, O. (2008) After the hotspots are gone: Land use history and grassland plant species diversity in a strongly transformed agricultural landscape. Applied Vegetation Science, 11, 365–374.

Foody, G.M. (2002) Status of land cover classification accuracy assessment. Remote Sensing of Environment, 80, 185–201.

Gartz, M. (2015) Plantdiversitet på svenska slåtterängar : En GIS-analys med kulturella perspektiv. Bachelor thesis in Physical Geography at Stockholm University.

Greenberg, J.A. & Mattiuzzi, M. (2015) gdalUtils: Wrappers for the Geospatial Data Abstraction Library (GDAL) Utilities. R package version 2.0.1.7, url: http://CRAN.R-project.org/package=gdalUtils.

Hijmans, R.J. (2016) raster: Geographic Data Analysis and Modeling. R package version 2.5-8, url: http://CRAN.R-project.org/package=raster.

Hooftman, D.A.P. & Bullock, J.M. (2012) Mapping to inform conservation: A case study of changes in semi-natural habitats and their connectivity over 70 years. Biological Conservation, 145, 30–38.

Hooftman, D.A.P., Edwards, B. & Bullock, J.M. (2016) Reductions in connectivity and habitat quality drive local extinctions in a plant diversity hotspot. Ecography, 39, 583–592.

Jiang, M., Bullock, J.M. & Hooftman, D.A.P. (2013) Mapping ecosystem service and biodiversity changes over 70 years in a rural English county. Journal of Applied Ecology, 50, 841–850.

Newbold, T., Hudson, L.N., Arnell, A.P., Contu, S., Palma, A.D., Ferrier, S., Hill, S.L.L., Hoskins, A.J., Lysenko, I., Phillips, H.R.P., Burton, V.J., Chng, C.W.T., Emerson, S., Gao, D., Pask-Hale, G., Hutton, J., Jung, M., Sanchez-Ortiz, K., Simmons, B.I., Whitmee, S., Zhang, H., Scharlemann, J.P.W. & Purvis, A. (2016) Has land use pushed terrestrial biodiversity beyond the planetary boundary? A global assessment. Science, 353, 288–291.

R Development Core Team. (2015) R: A Language and Environment for Statistical Computing. R Foundation for Statistical Computing, Vienna.

Saar, L., Takkis, K., Pärtel, M. & Helm, A. (2012) Which plant traits predict species loss in calcareous grasslands with extinction debt? Diversity and Distributions, 18, 808–817.

Skånes, H.M. & Bunce, R.G.H. (1997) Directions of landscape change (1741–1993) in Virestad, Sweden — characterised by multivariate analysis. Landscape and Urban Planning, 38, 61–75.

Stamp, D.L. (1931) The Land Utilisation Survey of Britain. The Geographical Journal, 78, 40–47.

Swetnam, R.D. (2007) Rural land use in England and Wales between 1930 and 1998: Mapping trajectories of change with a high resolution spatio-temporal dataset. Landscape and Urban Planning, 81, 91–103.

Wickham, H. (2009) ggplot2 - Elegant Graphics for Data Analysis. Springer, New York.

Willcock, S., Phillips, O.L., Platts, P.J., Swetnam, R.D., Balmford, A., Burgess, N.D., Ahrends, A., Bayliss, J., Doggart, N., Doody, K., Fanning, E., Green, J.M.H., Hall, J., Howell, K.L., Lovett, J.C., Marchant, R., Marshall, A.R., Mbilinyi, B., Munishi, P.K.T., Owen, N., Topp-Jorgensen, E.J. & Lewis, S.L. (2016) Land cover change and carbon emissions over 100 years in an African biodiversity hotspot. Global Change Biology, 22, 2787–2800.

